# On a quantum-inspired kernel for classifying protein torsion angles

**DOI:** 10.1101/2025.08.05.668681

**Authors:** Ashar Malik, David Ascher

## Abstract

Algorithms grounded in quantum principles need to demonstrate they are at least as expressive as established classical methods before any hardware advantage can be sought. Here we demonstrate that a quantum-inspired kernel reaches the same accuracy and Matthews correlation coefficient as carefully tuned radial basis function and degree-2 polynomial support vector machines when tasked with separating geometric regions in Ramachandran space. On a rigorously curated, balanced dataset of 10,000 torsion angle pairs derived from DSSP-annotated residues, the model achieves 98% accuracy with a Matthews correlation coefficient of 0.96, comparable to the top-performing classical models. By achieving true predictive parity on a well-characterised structural benchmark, the quantum-inspired kernel establishes that quantum-informed similarity measures already match classical baselines, laying a firm groundwork for future quantum-native bioinformatics workflows once suitable hardware becomes available.

## Introduction

Quantum computing promises to transform the life sciences [1, 2], from first-principles molecular simulation to large-scale combinatorial optimisation and high-dimensional sampling [2]. Two hurdles, however, stand between promise and practice. First, mapping a biological problem onto quantum hardware requires non-trivial data encodings and objective reformulations, often ballooning circuit depth beyond any near-term benefit [3, 4, 5]. Second, today’s noisy-intermediate-scale quantum (NISQ) devices suffer from decoherence and gate infidelity that sharply limit useful circuit depth [6, 7, 8]. In parallel, *quantum-inspired* algorithms, classical methods that mirror the algebraic structure of shallow quantum circuits [6, 9] have gained traction as a hardware-agnostic way to explore quantum principles.

Among biological data science tasks, three categories are poised to benefit: *classification* (e.g., sequence analysis and genomics and, by extension, other areas such as fold prediction, structural phylogenetics etc.) [4, 8, 10, 11], *optimisation* (e.g., protein folding) [1, 2, 8, 12, 13], and *sampling* (conformational exploration of flexible macromolecules) [1, 6, 8]. Among these, classification is an attractive testbed because performance can be quantified empirically against well-calibrated classical baselines.

Here we examine a quantum-inspired classifier for the two backbone torsion angles, *phi* (*ϕ*) and *psi* (*ψ*), whose joint distribution forms the Ramachandran plot. Because both angles lie in the bounded interval [−180^*°*^, 180^*°*^] and decades of crystallography have mapped the geometry exhaustively, Ramachandran space provides an almost “closed-world” benchmark or in other words where differences in model accuracy arise primarily from the modelling approach and not data sparsity.

To enable this we use DSSP [14], which assigns per-residue secondary-structure states based on hydrogen-bond patterns and local geometry. In this study we do not classify DSSP states directly; instead, we estimate an empirical Ramachandran density from all DSSP-annotated residues, denoise it via thresholding and a simple post-processing step, and define the “allowed” support 𝒮 ⊂ [−180^*°*^, 180^*°*^]^2^ as the remaining region of detected density; its complement is then taken as “disallowed”. We then ask whether membership in this denoised support can be approximated using only the backbone torsion pairs (*ϕ, ψ*). Rather than reproducing the classic steric-clash map, this reframing casts the task as learning the geometric decision boundary in torsion space as a surrogate for DSSP’s structural judgement. Our goal is conceptual rather than accelerative: *can a quantum-inspired, fidelity-inspired dotproduct–squared kernel perform on par with strong classical kernels on this task?* A positive answer offers an early, hardware-agnostic gauge of the practical value that quantum principles may bring to biological machine-learning workflows.

In this work, each (*ϕ, ψ*) pair is mapped to a 12-dimensional (12D) trigonometric embedding that mirrors single-qubit rotation encodings. Similarity is then defined as the *square* of the inner product between embeddings, mimicking quantum-state fidelity as described in the Qiskit SDK [15]. The contribution thus lies in recasting a well-known kernel as a fidelity measure ready for shallow quantum hardware for a biological problem. We benchmark a support-vector machine (SVM) using this recast kernel against linear, radial-basis and polynomial SVMs across ten random train–test splits, reporting accuracy and Matthews correlation coefficient (MCC). While intentionally simple, the embedding → kernel → classifier pipeline generalises to other biological classification problems and can be ported directly to quantum circuits when suitable hardware becomes available.

## Methods

### Data Generation

To construct a well-characterised classification dataset for torsion-angle analysis, we derived per-residue (*ϕ, ψ*) torsion angles from experimentally determined protein structures in the RCSB Protein Data Bank (PDB) [16] (https://www.rcsb.org/). Backbone torsions were computed using DSSP, and a custom Python parser was used to extract (*ϕ, ψ*) for all annotated residues; torsions for terminal residues were excluded. We then estimated a single empirical Ramachandran density over (*ϕ, ψ*) using 1^*°*^ bins, applied a small probability threshold to remove low-support pixels, and used a connected-component filter to discard isolated artefacts. The surviving region defines the “allowed” support 𝒮 ⊂ [−180^*°*^, 180^*°*^]^2^; its complement in [−180^*°*^, 180^*°*^]^2^ is “disallowed”.

From these masks, we sampled *N* = 5000 points from *S* (label 1) and *N* = 5000 points from its complement (label 0) without replacement, yielding a balanced dataset of 10,000 labelled torsion pairs used for all downstream training and evaluation. Angles were converted to radians prior to computing trigonometric features.

### Quantum-Inspired Feature Expansion

Each (*ϕ, ψ*) pair is mapped to a 12-dimensional (12D) feature vector using a trigonometric embedding that mirrors the angle-encoding patterns of variational quantum circuits. Combinations of sine and cosine terms emulate amplitude and phase modulations in single-qubit rotations, providing a classical analogue of quantum state preparation.

The 12 features comprise the set {sin *ϕ*, cos *ϕ*, sin *ψ*, cos *ψ*, sin(*ϕ* + *ψ*), cos(*ϕ* + *ψ*), sin(*ϕ* − *ψ*), cos(*ϕ* − *ψ*), sin(*ϕ ψ*), cos(*ϕ ψ*), sin(2*ϕ*), cos(2*ψ*)*}*. Collectively, these features reproduce much of the expressive power of shallow, entangled quantum circuits while remaining computationally lightweight. This compact 12D set was chosen empirically for this task and fixed for all experiments to aid reproducibility.

### Kernels and Classification Models

To benchmark this approach, we trained four support-vector-machine (SVM) configurations, each evaluated over ten random train–test splits (five-fold internal cross-validation).

- **Linear kernel (12D)**. A standard linear kernel applied to the 12D trigonometric embedding to assess whether the feature map alone has the ability to produce a linearly separable representation.
- **Polynomial kernel (12D)**. A degree-2 polynomial kernel was applied to the 12D features to capture pairwise interactions among the embedded features. For this hyperparameters *C, γ*, and *coef*_0_ were tuned by grid search.
- **Radial-basis (RBF) kernel (2D)**. To establish a classical baseline in the native torsion space, an RBF kernel was applied directly to the raw (*ϕ, ψ*) angles (no embeddings).
- **Quantum-inspired kernel (12D)**. We define

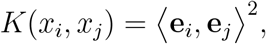

where **e**_*i*_ and **e**_*j*_ are the 12D trigonometric embeddings. Algebraically, *K* is a homogeneous degree-2 polynomial kernel operating on our embedding. We describe it as “fidelity-inspired” to signal its connection to overlap-based similarities used in shallow quantum circuits, while keeping the implementation purely classical to address our central question: *do quantum-inspired kernels work well on biological data?* A fidelity-exact variant can be obtained by unit-normalizing embeddings (or normalizing the Gram matrix), but this was not required for the present benchmark. The kernel matrix was precomputed and passed to `SVC` via `kernel=‘precomputed’`.

### Web Application Deployment

To make the models accessible and interpretable, we developed a web application using the Flask framework. Users can upload PDB files or retrieve structures directly from the RCSB PDB. Backbone torsion angles are computed server-side, using DSSP, and classified into allowed or disallowed regions using both the RBF (baseline) and quantum-inspired SVM models. The application provides an interactive Mol* viewer [17] for 3D structure visualisation, and uses RCSB-Saguaro [18] as a feature viewer that overlays DSSP-derived ground truth with model predictions from both kernels. Probability maps for each model are displayed over the torsion-angle space. Users can probe decision boundaries via an interactive sampler that draws (*ϕ, ψ*) pairs from the empirical 2D Ramachandran density (thresholded and morphologically cleaned) and evaluates them with both classifiers; the sampled points and predictions can be downloaded. An API is also available to enable programmatic access.

## Results

### Torsion-Space Data Coverage

DSSP-derived (*ϕ, ψ*) angles for all RCSB PDB structures were first grouped into *allowed* and *disallowed* regions (Supplementary Fig. S1). Sparse, isolated pixels were removed with a connected-component filter to obtain a contiguous representation of conformational space. From the cleaned masks we then sampled an equal number of points from each class, yielding a balanced dataset of 10,000 residues (5,000 allowed and 5,000 disallowed; Supplementary Fig. S2).

### Classification Performance

Table 1 summarises the mean test accuracy and MCC across ten random train–test splits. The linear kernel applied to the 12D embedding performs relatively poorly, confirming that the feature map alone is not linearly well separable. Introducing non-linearity with an RBF kernel in raw torsion space raises accuracy to roughly 96%. Both the degree-2 polynomial kernel on the 12D features and the quantum-inspired kernel reach the highest scores, with statistically indistinguishable metrics.

**Table 1:** Mean (*±* SD) accuracy and MCC over ten seeds. Bold values indicate the two top-performing kernels.

| Kernel / Feature set | Accuracy (%) | MCC |
| --- | --- | --- |
| Linear (12D) | $76.85 \pm 0.92$ | $0.537 \pm 0.018$ |
| RBF (2D) | $96.27 \pm 0.40$ | $0.926 \pm 0.008$ |
| **Poly-2 (12D)** | **$97.93 \pm 0.28$** | **$0.959 \pm 0.005$** |
| **Quantum-inspired (12D)** | **$97.97 \pm 0.22$** | **$0.960 \pm 0.004$** |

## Discussion

Contemporary quantum processors remain in the noisy-intermediate-scale (NISQ) era, where decoherence, gate infidelity and limited qubit counts constrain circuit depth and computational accuracy. Classical simulators offer a valuable testbed, yet their exponential memory demands restrict them to toy problems. In this context, *quantum-inspired* algorithms provide a pragmatic bridge: they allow exploration of the algebraic structure of shallow quantum circuits while running on conventional hardware, allowing workflows to be “quantum-ready” before fault-tolerant devices arrive.

Throughout the paper we use “allowed” and “disallowed” in an operational sense: (*ϕ, ψ*) pairs inside the denoised empirical support are labelled allowed, and those outside are labelled disallowed. In other words, the classifier designed in this work does not reproduce the classic steric-clash Ramachandran map; instead it asks whether two backbone torsion angles alone can approximate the hydrogen-bond and geometry test carried out by DSSP. Achieving almost 98% accuracy therefore shows that torsion angles (*ϕ, ψ*) capture a substantial fraction of the signal DSSP uses to infer the state of a residue.

The choice of simple and fully interpretable dataset, Ramachandran torsion angles, is thus deliberate, and allows empirical examination of the value of a fidelity-motivated similarity measure without confounding data issues. Each torsion-angle (*ϕ, ψ*) pair was embedded in a 12D trigonometric space that mirrors single-qubit rotation encodings, and similarity was taken as the *square* of the inner product between embeddings. Algebraically, this kernel is a degree-2 polynomial; conceptually, it is a fidelity-inspired dot-product–squared similarity on circuit-like features. With this reinterpretation in place, the resulting SVM matched the accuracy and MCC of strong classical baselines (RBF and poly-2) while outperforming a plain linear kernel applied to the same features. Because timing differences depend strongly on solver choice, SVC for the polynomial model versus a precomputed Gram matrix for the fidelity-inspired kernel, we emphasise predictive parity rather than speed-up: on a noise-free, well-characterised dataset, a quantum-inspired kernel is on par with established methods.

To make the approach accessible, we have released a publicly available web server focused on Ramachandran classification. The site lets users upload or fetch protein structures, visualise DSSP ground truth, and compare per-residue predictions from an RBF and the quantum-inspired kernel side by side. By linking model output directly to experimental labels, the web application provides an interpretable dashboard from which to assess quantum and classical decision boundaries in a familiar biological setting.

Many biological classification tasks are naturally binary (e.g., benign versus pathogenic variants, active versus inactive compounds, folded versus misfolded proteins). The present study shows that quantum-inspired kernels are easy to implement, require no specialised hardware, and integrate seamlessly into existing machine-learning pipelines. We also speculate that this benchmark has the potential to be used as a framework for developing and testing noise correction techniques, which will be critical for achieving reliable performance on true quantum hardware. As quantum technology matures, workflows already cast in fidelity-like kernels will be primed for direct deployment on fault-tolerant devices, potentially unlocking advantages that remain inaccessible to today’s purely classical approaches.

## Conclusion

Quantum-inspired kernels provide a practical avenue for testing quantum principles on today’s classical hardware. Here we showed that our quantum-inspired kernel, which is algebraically a degree-2 polynomial on our embedding, matches the accuracy and MCC of strong classical baselines when classifying allowed versus disallowed regions in Ramachandran space. To help the community explore this perspective, we have released an interactive web server that lets users upload or fetch protein structures, visualise DSSP ground truth, and compare predictions from a conventional RBF SVM with those of the quantum-inspired model side by side. This interface makes the kernel’s behaviour transparent on real structural data.

Demonstrating predictive parity on a clean, well-understood benchmark and providing an open toolset paves the way for quantum-inspired kernels to be applied to broader biological tasks, such as benign versus pathogenic variant classification or active versus inactive ligand prediction well before large-scale quantum computers become commonplace.

## Availability

The interactive web server is freely accessible at https://biosig.lab.uq.edu.au/q_torsion.

## References

[1] Yudong Cao, Jonathan Romero, Jonathan P Olson, Matthias Degroote, Peter D Johnson, Mária Kieferová, Ian D Kivlichan, Tim Menke, Borja Peropadre, Nicolas PD Sawaya, et al. Quantum chemistry in the age of quantum computing. Chemical reviews, 119(19):10856–10915, 2019.

[2] Laura Marchetti, Riccardo Nifosí, Pier Luigi Martelli, Eleonora Da Pozzo, Valentina Cappello, Francesco Banterle, Maria Letizia Trincavelli, Claudia Martini, and Massimo D’Elia. Quantum computing algorithms: getting closer to critical problems in computational biology. Briefings in Bioinformatics, 23(6):bbac437, 2022.

[3] Jarrod R McClean, Sergio Boixo, Vadim N Smelyanskiy, Ryan Babbush, and Hartmut Neven. Barren plateaus in quantum neural network training landscapes. Nature communications, 9(1):4812, 2018.

[4] Vojtěch Havlíček, Antonio D Córcoles, Kristan Temme, Aram W Harrow, Abhinav Kandala, Jerry M Chow, and Jay M Gambetta. Supervised learning with quantum-enhanced feature spaces. Nature, 567(7747):209–212, 2019.

[5] Maria Schuld, Ryan Sweke, and Johannes Jakob Meyer. Effect of data encoding on the expressive power of variational quantum-machine-learning models. Physical Review A, 103(3):032430, 2021.

[6] John Preskill. Quantum computing in the nisq era and beyond. Quantum, 2:79, 2018.

[7] Morten Kjaergaard, Mollie E Schwartz, Jochen Braumüller, Philip Krantz, Joel I-J Wang, Simon Gustavsson, and William D Oliver. Superconducting qubits: Current state of play. Annual Review of Condensed Matter Physics, 11(1):369–395, 2020.

[8] Kishor Bharti, Alba Cervera-Lierta, Thi Ha Kyaw, Tobias Haug, Sumner Alperin-Lea, Abhinav Anand, Matthias Degroote, Hermanni Heimonen, Jakob S Kottmann, Tim Menke, et al. Noisy intermediate-scale quantum algorithms. Reviews of Modern Physics, 94(1):015004, 2022.

[9] Ewin Tang. A quantum-inspired classical algorithm for recommendation systems. In Proceedings of the 51st annual ACM SIGACT symposium on theory of computing, pages 217–228, 2019.

[10] Vedran Dunjko and Hans J Briegel. Machine learning & artificial intelligence in the quantum domain: a review of recent progress. Reports on Progress in Physics, 81(7):074001, 2018.

[11] Prashant S Emani, Jonathan Warrell, Alan Anticevic, Stefan Bekiranov, Michael Gandal, Michael J McConnell, Guillermo Sapiro, Alán Aspuru-Guzik, Justin T Baker, Matteo Bastiani, et al. Quantum computing at the frontiers of biological sciences. Nature Methods, 18(7):701–709, 2021.

[12] Anton Robert, Panagiotis Kl Barkoutsos, Stefan Woerner, and Ivano Tavernelli. Resource-efficient quantum algorithm for protein folding. npj Quantum Information, 7(1):38, 2021.

[13] Ashar J Malik and Chandra S Verma. On quantum computing and geometry optimization. bioRxiv, pages 2023–03, 2023.

[14] Wolfgang Kabsch and Christian Sander. Dictionary of protein secondary structure: pattern recognition of hydrogen-bonded and geometrical features. Biopolymers: Original Research on Biomolecules, 22(12):2577–2637, 1983.

[15] Ali Javadi-Abhari, Matthew Treinish, Kevin Krsulich, Christopher J Wood, Jake Lishman, Julien Gacon, Simon Martiel, Paul D Nation, Lev S Bishop, Andrew W Cross, et al. Quantum computing with qiskit. arXiv preprint 2405.08810, 2024.

[16] Helen M Berman, John Westbrook, Zukang Feng, Gary Gilliland, Talapady N Bhat, Helge Weissig, Ilya N Shindyalov, and Philip E Bourne. The protein data bank. Nucleic acids research, 28(1):235–242, 2000.

[17] David Sehnal, Sebastian Bittrich, Mandar Deshpande, Radka Svobodová, Karel Berka, Václav Bazgier, Sameer Velankar, Stephen K Burley, Jaroslav Koča, and Alexander S Rose. Mol* viewer: modern web app for 3d visualization and analysis of large biomolecular structures. Nucleic acids research, 49(W1):W431–W437, 2021.

[18] Joan Segura, Yana Rose, John Westbrook, Stephen K Burley, and Jose M Duarte. Rcsb protein data bank 1d tools and services. Bioinformatics, 36(22-23):5526–5527, 2020.

