## Supplementary Fig for "On a quantum-inspired kernel for classifying protein torsion angles"

### Supplementary Figures

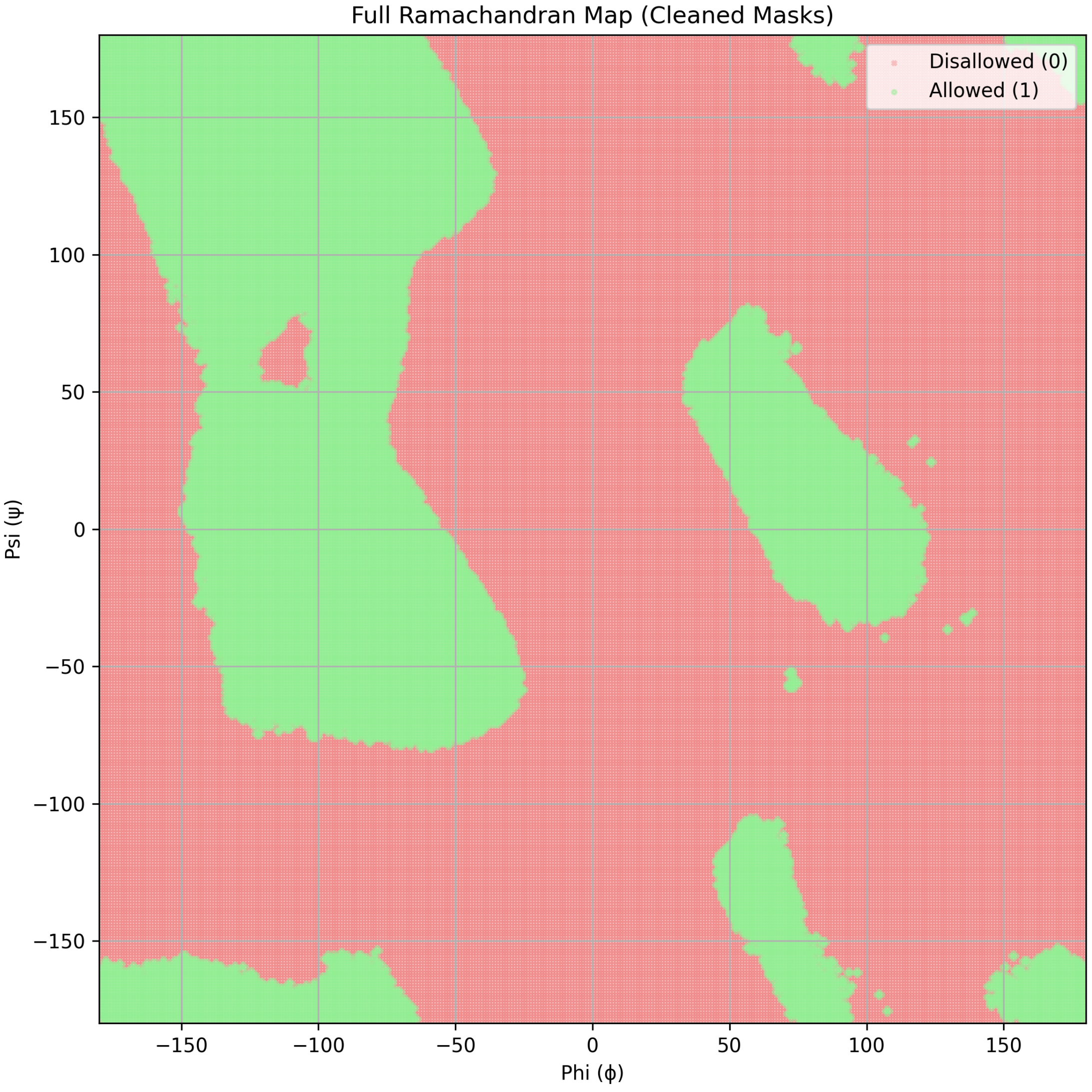


Figure S1: Cleaned Ramachandran density map after removing sparse pixels. Allowed regions are shown in green and disallowed regions in red.

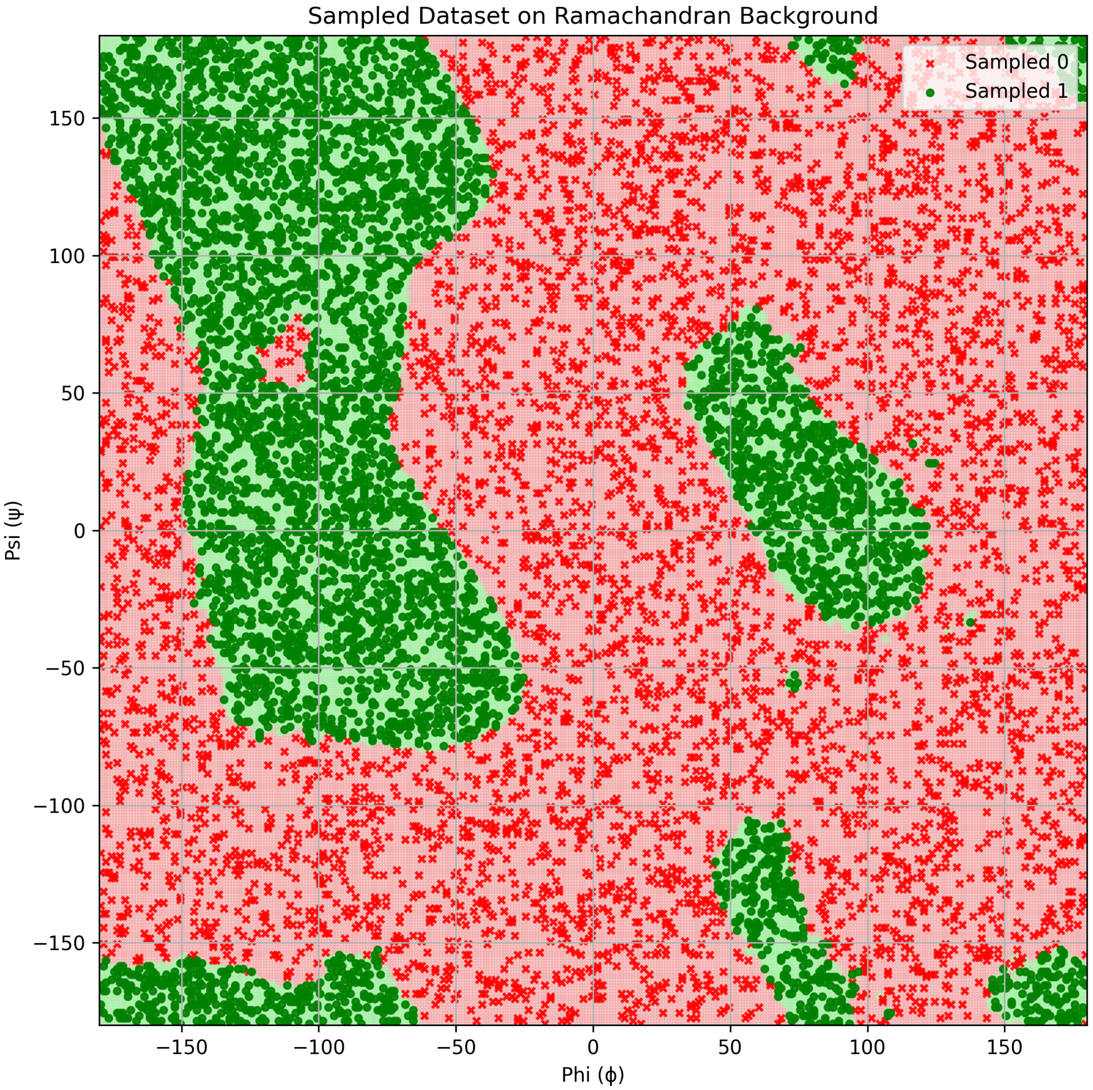


Figure S2: Balanced training dataset ($N = 10{,}000$) sampled from the cleaned Ramachandran map. Red crosses represent disallowed (Sampled 0) and green circles represent allowed (Sampled 1) torsion-angle configurations.
